# Effect of SARS-CoV-2 spike mutations on its activation by TMPRSS2 and TMPRSS13

**DOI:** 10.1101/2022.01.26.477969

**Authors:** Annelies Stevaert, Ria Van Berwaer, Valerie Raeymaekers, Manon Laporte, Lieve Naesens

## Abstract

The continuous emergence of new SARS-CoV-2 variants urges better understanding of the functional motifs in the spike (S) protein and their tolerance towards mutations. We here focus on the S2’ motif which, during virus entry, requires cleavage by a cell surface protease to release the fusion peptide. Though belonging to an immunogenic region, the SARS-CoV-2 S2’ motif (811-KPSKR-815) has shown hardly any variation, with its three basic (K/R) residues being >99.99% conserved thus far. By creating a series of mutant S-pseudotyped viruses, we show that K_814_, which precedes the scissile R_815_ residue, is dispensable for SARS-CoV-2 spike activation by TMPRSS2 but not TMPRSS13. The latter protease lost its activity towards SARS-CoV-2 S when the S2’ motif was swapped with that of the low pathogenic 229E coronavirus (685-RVAGR-689) and also the reverse effect was seen. This swap had no impact on TMPRSS2 activation. Also in the MERS-CoV spike, introducing a dibasic scissile motif was fully accepted by TMPRSS13 but less so by TMPRSS2. Our findings are the first to demonstrate which S2’ residues are important for SARS-CoV-2 spike activation by these two airway proteases, with TMPRSS13 exhibiting higher preference for K/R rich motifs than TMPRSS2. This preemptive insight can help to estimate the impact of S2’ motif changes as they may appear in new SARS-CoV-2 variants.

**IMPORTANCE:** Since the start of the COVID-19 pandemic, SARS-CoV-2 is undergoing worldwide selection with frequent appearance of new variants. The surveillance would benefit from proactive characterization of the functional motifs in the spike protein, the most variable viral factor. This is linked to immune evasion but also influences spike functioning in a direct manner. Remarkably, though located in a strong immunogenic region, the S2’ cleavage motif has, thus far, remained highly conserved. This suggests that its amino acid sequence is critical for spike activation by airway proteases. To investigate this, we assessed which S2’ site mutations affect processing by TMPRSS2 and TMPRSS13, two main activators of the SARS-CoV-2 spike. Being the first in its kind, our study will help to assess the biological impact of S2’ site variations as soon as they are detected during variant surveillance.

---

Since the start of the COVID-19 pandemic, SARS-CoV-2 is undergoing worldwide selection with frequent appearance of new variants. The variability in the viral spike (S) antigen is linked to immune evasion but also affects the functioning of S in virus replication and transmission. For instance, substitution D614G, which arose in March 2020 to soon become dominant, increases S protein stability (1, 2) and virus transmission (3, 4). Mutation P681R, present in the delta and kappa variants and located adjacent to the S1/S2 furin recognition motif (RRAR), enhances spike fusogenicity and virus pathogenicity in hamsters (5). The reverse is seen for the omicron variant (6, 7). Knowing which residues in the functional spike motifs are essential or not, can help to assess the impact of new variations as they emerge. In this report, we focus on the S2’ motif.

Cleavage of the SARS-CoV-2 S1/S2 site facilitates cleavage at the S2’ site, a process essential to release the fusion peptide. Two efficient S2’ activators are TMPRSS2 (8) and TMPRSS13 (2, 9, 10) which are both expressed in respiratory tissue (11). In a SARS-CoV-2 mouse model, TMPRSS2 knockout resulted in less lung pathology and lower virus titers, although the virus was still able to replicate in the lungs (12). In human airway-derived Calu-3 cells, knockdown of TMPRSS2 reduced virus replication dramatically (2, 13), yet also TMPRSS13 knockdown had significant effect (2). The activating role of TMPRSS2, but not that of TMPRSS13, was confirmed in human airway organoids (14). Hence, the relative contribution of these two proteases in SARS-CoV-2 infection remains unclear.

## Low variability and K/R abundance in the S2’ site

Within the SARS-CoV-2 S2’ motif (811-KPSKR-815), the scissile R_815_ residue is flanked by a second basic (K_814_) residue. The motif lies in a strong epitope (15, 16) for which the antibody titers seem correlated with COVID-19 disease severity (17). Despite this high immunogenicity, the S2’ motif exhibits strikingly low variability. When we analyzed the ~6.7 million spike sequences in the GISAID database (Fig. 1A), all three basic residues in the S2’ motif (= K_811_, K_814_ and R_815_) proved highly dominant, being present in >99.99% of the sequences. This is less surprising for R_815_ (in only a few cases substituted by K), the presumed scissile residue (18). P_812_ and S_813_ are somewhat more tolerant towards variation, consistent with the presence of T_813_ in the SARS-CoV spike (19).

**FIGURE 1.**
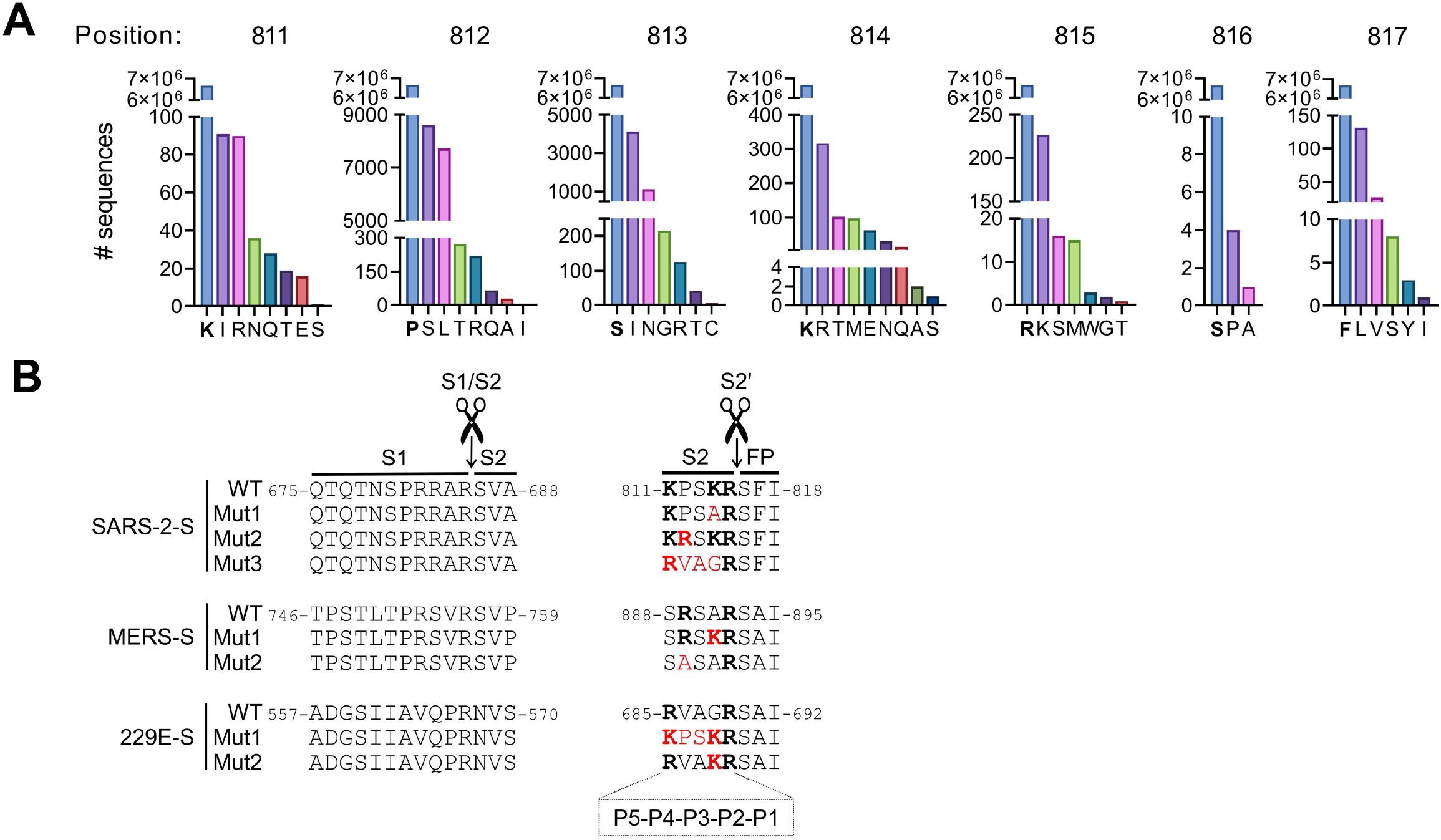
S2’ site variation in circulating SARS-CoV-2 isolates and mutations introduced in this study. **A.** Variation analysis of the S2’ site was performed on 6,679,700 SARS-2-S sequences downloaded from the GISAID database. **B.** Partial alignment of wild-type (WT) SARS-2-S, MERS-S and 229E-S, and the S2’ mutants created in this study. In red: amino acid changes versus WT; in bold: basic (i.e. R or K) residues.

## Mutations at the S2’ motif do not affect spike expression or S1/S2 cleavage

The high conservation of the SARS-CoV-2 KxxKR motif suggests that its sequence is essential for spike activation. Strikingly, a dibasic scissile motif is missing in all five endemic less pathogenic human coronaviruses (HCoVs) including HCoV-229E (19), while MERS-CoV bears an xRxxR motif (Fig. 1B). Hence, we designed a series of S2’-mutated S proteins (abbreviated SARS-2-S, MERS-S and 229E-S; Fig. 1B) in which we swapped the motifs from SARS-2-S and 229E-S or introduced or removed a Lys (K) at P2 or an Arg (R) at P4 (since MERS-S contains a basic residue at P4 but not P5). S-bearing MLV-pseudoparticles were produced in HEK293T cells, pelleted down and analyzed for spike levels and cleavage by western blot (Fig. 2A). All S2’-mutant pseudovirions contained similar S protein levels as the respective WT (Fig. 2B). For the SARS-2-S- and MERS-S-pseudoparticles, efficient spike cleavage was seen for all S2’mutants. Even a change of 4 out of 5 residues (= SARS-2-S-Mut3) had no effect on spike expression or S1/S2 cleavage. For 229E-S, the WT and mutants contained similar levels of several cleavage products.

**FIGURE 2.**
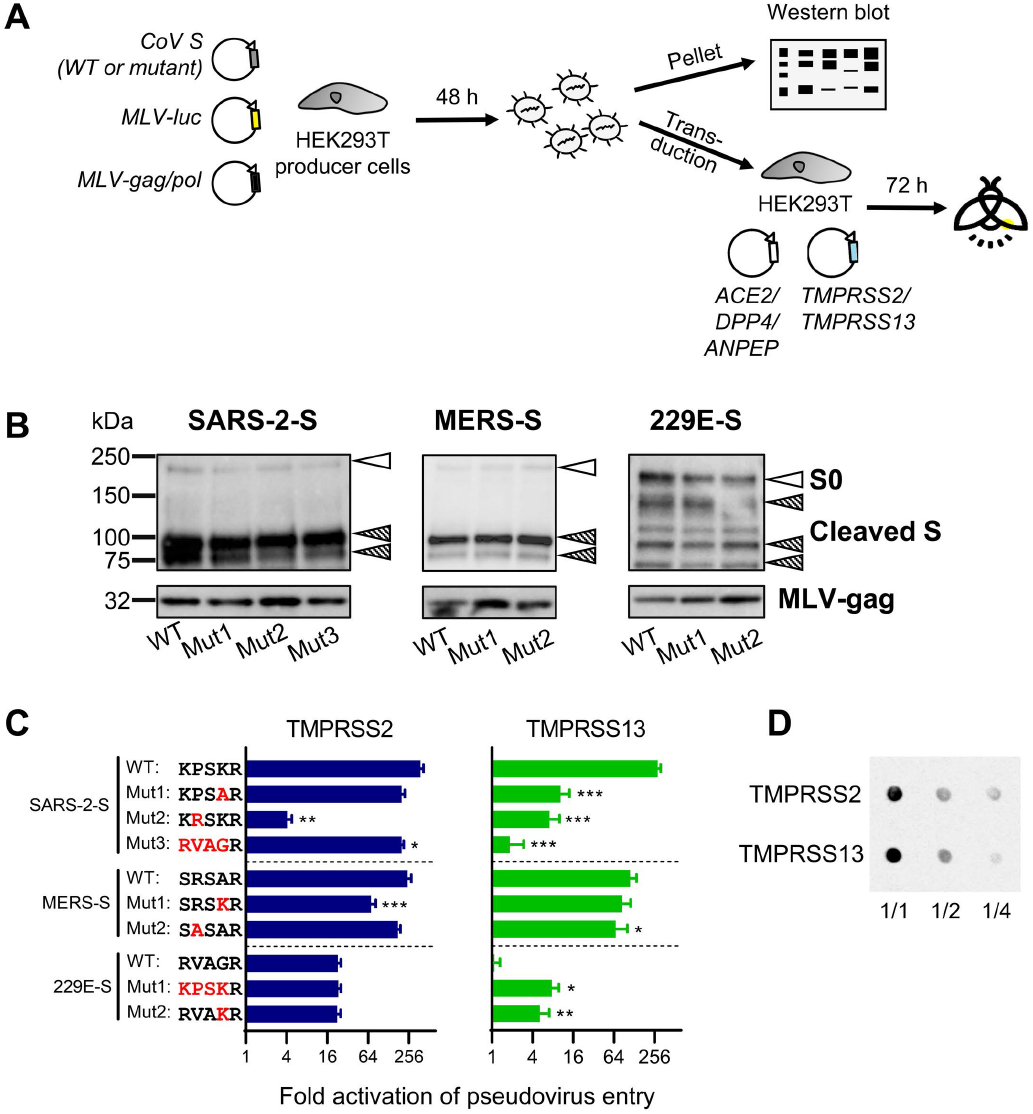
Impact of S2’ site changes on virus entry activation by TMPRSS2 and TMPRSS13. **A.** Experimental set-up. **B.** Spike protein levels and cleavage state in S-pseudotyped virions. The particles were produced in HEK293T cells, pelleted and submitted to western blot analysis with anti-V5 antibody recognizing the SO and S2 forms (top) and anti-MLV-gag antibody (bottom). Open arrows: full-length (SO) spike; dashed arrows: cleaved forms resulting from S1/S2 processing (SARS-2-S and MERS-S) or less specific cleavage (229E-S). **C.** Entry of WT and mutant S-pseudotyped viruses into HEK293T cells transfected for TMPRSS2 or TMPRSS13. The bars show the factor activation, i.e. luminescence signal relative to the condition receiving empty instead of protease-plasmid. Pooled data from 3 experiments, each performed in 6 replicates. *, P ≤0.05; **, P <0.01; ***, P ≤0.001 (nested t-test, two-tailed). **D.** Dot blot analysis of TMPRSS2 and TMPRSS13 expression in the transfected HEK293T cells. The cell extracts were spotted undiluted (1/1) or diluted 1/2 or 1/4, and detection was done with anti-flag tag antibody.

## SARS-2-S containing the S2’ motif of 229E-S loses TMPRSS13 cleavability

We next conducted an entry assay to measure S2’ activation by TMPRSS2 and TMPRSS13. The S-pseudoviruses were transduced into HEK293T cells which were transfected one day earlier with TMPRSS2-, TMPRSS13- or empty plasmid, plus the cognate virus receptor (Fig. 2A). Transduction was performed in the presence of E64d to shut off the endosomal route in which S is activated by cathepsin B/L instead of cell surface proteases (20).

As expected (2, 8, 21), TMPRSS2 proved an efficient activator of the three CoV spike proteins, giving an increase in S-driven pseudovirus entry (compared to cells transfected with empty plasmid) of 378, 244 and 23 for the WT forms of SARS-2-S, MERS-S and 229E-S, respectively (Fig. 2C). The lower value for 229E-S is explained by lower levels of functional S in these pseudovirions, due to unspecific cleavage (see above). 229E-S was not activated by TMPRSS13, in sharp contrast with efficient usage of this protease by SARS-2-S and MERS-S. Regarding the S2’ mutants, neither was severely affected for TMPRSS2 usage. The only exception was SARS-2-S-Mut2 showing that TMPRSS2-driven entry was 92-fold reduced when the P at P4 was substituted by R. Removing the basic charge at P2 (= K to A; Mut1) or swapping the motif of SARS-2-S for that of 229E-S (Mut3) gave only 2-fold reduction in activation by TMPRSS2. In stark contrast, these mutations had significantly (P < 0.001) negative impact on TMPRSS13-driven entry, the fold reduction versus SARS-2-S WT being 27- (Mut1), 40- (Mut2) and as high as 155-fold for Mut3. Hence, introducing the 229E-S2’ motif into SARS-2-S abolished the capacity of TMPRSS13 to activate this spike. Inversely, 229E-S gained the ability to use TMPRSS13 for entry activation, when bearing the SARS-2-S2’ motif (Mut1; 7.1-fold) or its K at P2 (Mut2; 4.7-fold).

For MERS-S, activation by TMPRSS2 was significantly yet only 3.4-fold reduced when the motif contained a third basic residue (Mut1). Substituting the R at P4 (Mut2) had no (TMPRSS2) or only minor (TMPRSS13) influence.

During the above experiments, equal amounts of TMPRSS2- and TMPRSS13-plasmid DNA were applied for HEK293T cell transfection. Dot blot analysis demonstrated that this generated comparable protein levels for both proteases (Fig. 2D). This excluded the possibility that low expression of TMPRSS13 might be the reason for poor activation of some pseudoviruses.

## Discussion

This study anticipates on the relevance of S2’ site changes as they may appear in new SARS-CoV-2 variants. Since the S2’ motif lies in a strong epitope (15, 17), the virus might acquire mutations in this region to evade immunity. The fact that such variations are, thus far, rarely detected could mean that the sequence of this K/R rich motif is crucial for SARS-2-S activation by cell surface proteases. Our study is the first to address this topic. Clearly, residue K_814_ is important for TMPRSS13 but not TMPRSS2. This basic P2 residue differentiates SARS-2-S from 229E-S (and the spikes of all other low pathogenic HCoVs), which otherwise share a basic P5 residue. Making the motif even more basic by addition of an R at P4, causes a more dramatic reduction for TMPRSS2 than TMPRSS13. This aligns with data that basic residues at P2-P3-P4 are not favored by TMPRSS2 (22). Reciprocally, the preference of TMPRSS13 for a basic motif concurs with its activity on multibasic hemagglutinins of highly pathogenic avian influenza viruses (23). Combined with the presence of TMPRSS13 in human lung tissue (24), this raises the hypothesis that loss of TMPRSS13 cleavability, e.g. by mutation of K_814_, might reduce replication of SARS-CoV-2 in the lungs. Though limited to a few mutations, our findings will help to estimate the impact of S2’ site changes observed during variant surveillance.

## Experimental procedures

In this study, the WT spike sequences correspond to the published sequences with accession numbers: YP_009724390.1 (SARS-2-S; early pandemic D614 variant); YP_009047204.1 (MERS-S) and NP_073551.1 (229E-S). A detailed description of the methods under (i) to (iii) can be found elsewhere (2). **(i) Pseudovirus production.** Briefly, the plasmids encoding the WT spikes (with C-terminal V5-tag) were first submitted to site-directed mutagenesis. To produce S-pseudotyped murine leukemia virus (MLV) particles, the S-plasmids were combined with the MLV gag-pol and firefly luciferase reporter plasmids and co-transfected into HEK293T cells. The pseudovirus-containing supernatants were collected after three days incubation at 33°C (SARS-2-S and 229E-S) or 37°C (MERS-S). **(ii) Effect of**

## TMPRSS2 and TMPRSS13 on pseudovirus entry

HEK293T cells were transfected with TMPRSS2-, TMPRSS13- or empty plasmid (11) combined with a plasmid encoding the suitable coronavirus receptor, i.e. angiotensin-converting enzyme 2 (ACE2; for SARS-2-S); dipeptidyl peptidase-4 (DPP4; for MERS-S) or aminopeptidase N (APN; for 229E-S). One day later, the cells were pre-incubated for 2 h with cathepsin inhibitor E64d, after which pseudovirus was added and allowed to enter for 2 h. After replacing the medium, the cells were incubated for three days at 33°C, to then measure the luciferase signal in a plate luminometer. **(iii) Western blot analysis of spike expression and cleavage.** After pseudovirus production [see (i)], the particles were pelleted from the supernatants by centrifugation on a sucrose cushion, after which they were mixed with RIPA buffer and submitted to denaturating PAGE. After blotting, the membranes were stained with anti-V5 and anti-MLV gag p30 antibodies combined with HRP-linked secondary antibody. **(iv) Dot blot detection of TMPRSS2 and TMPRSS13.** Two days after transfection with protease expression plasmid, extracts of the HEK293T cells were prepared in RIPA buffer, then spotted on nitrocellulose membranes in undiluted, or 1/2 and 1/4 diluted form. The membranes were stained with anti-flag antibody recognizing the N-terminal flag tag in the expressed TMPRSS2 and TMPSS13. All details can be found in a previous report (11). **(v) Variation analysis.** 6,679,700 spike sequences were downloaded from the GISAID database (25) on 04/01/2022 and variation around the S2’ site was analyzed.

## ACKNOWLEDGMENTS

The authors thank S. Pöhlmann and M. Hoffmann for the kind gift of plasmid materials.

